# Alcohol-sourced acetate impairs T cell function by promoting cortactin acetylation

**DOI:** 10.1101/2022.03.08.483253

**Authors:** Vugar Azizov, Michael Frech, Michel Hübner, Jörg Hofmann, Marketa Kubankova, Dennis Lapuente, Matthias Tenbusch, Jochen Guck, Georg Schett, Mario M. Zaiss

## Abstract

Alcohol is among the most widely consumed dietary substances. Excessive alcohol consumption damages the liver, heart and brain. Alcohol also has strong immunoregulatory properties. Here we report how alcohol impairs T cell function via acetylation of cortactin, a protein that binds filamentous actin and facilitates branching. Upon alcohol consumption, acetate, the metabolite of alcohol, accumulates in lymphoid organs. T cells exposed to acetate, exhibit increased acetylation of cortactin. Acetylation of cortactin inhibits filamentous actin binding and hence reduces T cell migration, immune synapse formation and activation. While mutated, acetylation-resistant cortactin rescued the acetate-induced inhibition of T cell migration, primary mouse cortactin knock-out T cells exhibited impaired migration. Furthermore, acetate-induced cytoskeletal changes effectively inhibited activation, proliferation, and immune synapse formation in T cells in vitro and in vivo in an influenza infection model in mice. Together these findings reveal cortactin as a possible target for mitigation of T cell driven autoimmune diseases.

## Introduction

For appropriate immune response to occur, immune cells such as T cells, must migrate within and outside of lymphoid organs ^1–4^. The cytoskeleton plays a vital role during T cell migration, scanning for antigens, and cell activation ^5–7^. To migrate, the cytoskeletal structures must remain highly dynamic ^3^. Failure in cytoskeletal machinery leads to immunodeficiencies due to defects in immune synapse formation, chemotaxis, and migration ^8^. Modulation of the components of cytoskeleton via post-translational modifications such as acetylation provides further fine tuning to fit cell’s demands ^9^. Histone deacetylases (HDAC) reverse acetylation of variety of proteins and have been shown to impact immunomodulatory effects of T cells ^10^. HDAC6 primarily deacetylates cytoskeletal proteins that has been shown to be required for proper immune synapse formation and migration ^11–16^. For example, pharmacological inhibition of HDAC6 in a preclinical model of systemic lupus erythematosus (SLE) reduced infiltrating T follicular helper (T_FH_) cells into germinal centers (GCs) with significant consequences on autoantibody titers ^13,14^. Similarly, we have previously shown that increased exposure to alcohol and alcohol’s main metabolite, acetate, affect T_FH_ cell responses in the collagen induced arthritis (CIA) – a mouse model of inflammatory arthritis ^17^. Mice exposed to either alcohol-sourced or directly supplemented acetate exhibited reduced T_FH_ cell infiltration of B cell follicles and destabilized T_FH_-B cell conjugate formation both *in vivo* and *in vitro* ^17^. Consequently, we observed acetate’s suppression to be specific to T cell dependent humoral responses both in autoimmune and vaccination mouse models ^17^. Inhibition of HDAC6 in the context of inflammatory arthritis has been shown to reduce disease severity to levels comparable to dexamethasone treatment ^12^. Here we hypothesized that acetate’s inhibitory effect could be due to increased acetylation of the cytoskeletal proteins ultimately affecting T cell migration and function. Our hypothesis could explain alcohol’s double-edged sword effect in regard to benefits in autoimmunity and damages to health in general ^18,19^. Alcohol consumption has correlated with decreased severity in rheumatoid arthritis, type 1 diabetes, SLE, and multiple sclerosis in humans as well as in disease mouse models ^19^. Upon consumption, alcohol is rapidly metabolized to acetate contributing to increased serum concentrations ^20^. Acetate permeates cells where it is converted to acetyl-Co-enzyme A (acetyl-CoA) and used as a donor for protein acetylation ^21,22^. Indeed, increased levels of intracellular acetyl-CoA were linked with significantly higher protein acetylation ^23^. In 2019, Mews and colleagues showed that alcohol consumption can quickly lead to histone acetylation in the brain ^24^. Considering acetate’s potential to increase acetylation of intracellular proteins, we set out to investigate whether there is a link between acetate exposure and mitigation of cytoskeletal dynamics in T cells.

## Results

### Upon alcohol consumption acetate accumulates in lymphoid organs

Building upon our published results, we ventured to study whether the decreased infiltration of GCs by T and B cells along with reduced Tfh-B cell contacts was due to cytoskeletal deficiencies following alcohol-sourced acetate exposure ^17^. First, we set to quantify the exposure of T cells to acetate upon alcohol consumption by measuring acetate levels in alcohol-impacted and immune relevant lymphoid tissues such as the liver, inguinal lymph node (iLN), and spleen of alcohol-fed mice. We found that acetate indeed accumulates in these organs, especially in lymphoid organs reaching concentrations surpassing 5 mM (Figures 1A-C). Next, we quantified whether exposure to 5 mM acetate concentration results in incorporation to cellular metabolism. Previously it was shown that increased *in vivo* ethanol concentration can lead to an increase in intracellular acetate and citrate ^24^. In an *in vitro* treatment of mouse naïve CD4^+^ CD25^-^ CD44^low^ CD62L^high^ T cells, hereafter naïve CD4^+^ T cells, with ^13^C labelled acetate, we were able to confirm an increase in ^13^C containing citrate starting at exposure to 2 mM acetate concentrations (Figure 1D). In a subsequent quantification of histone 3 (H3) acetylation levels in the same CD4^+^ T cells, we observed increased H3 acetylated at lysine 27 (Figure 1E). Together, these findings confirm increased exposure of T cells to acetate, leading to increased protein acetylation within cells.

**Figure 1.**
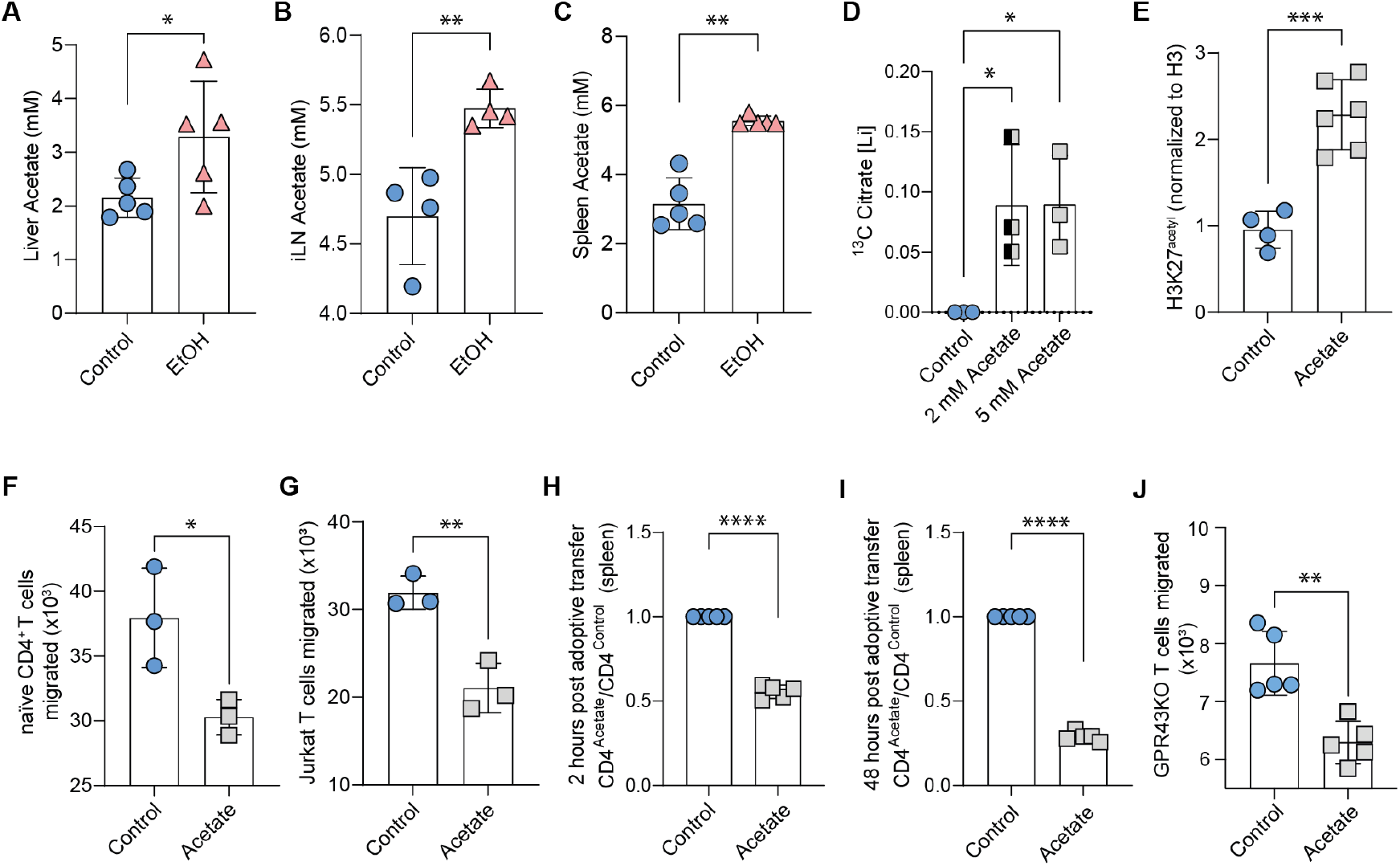
Acetate accumulates in lymphoid organs and impairs T cells migration capacity. GC/MS quantification of acetate in (**A**) livers, (**B**) inguinal lymph nodes (iLN), and (**C**) spleens of alcohol-fed mice compared with water-fed control mice, expressed in mM concentration (individual data point represents one mouse). (**D**) Mouse CD4^+^ T cells treated *in vitro* with 5 mM ^13^C labelled acetate exhibit increased ^13^C labelled citrate, quantified by GC-MS. (**E**) Quantification of western blot analysis of the level of acetylated histone 3 at lysine 27 of mouse CD4^+^ T cells treated *in vitro* with 5 mM acetate. (**F**) Numbers of migrated mouse naïve CD4^+^ T cells and (**G**) Jurkat T cells *in vitro* trans-well assay, counted by flow cytometer. (**H**) The ratio of acetate to control treated mouse CD4^+^ T cells found in the spleens of recipient mice in an adoptive transfer experiment at 2 hours and (**I**) 48 hours post transfer (each point represents one mouse, n = 5). (**J**) Numbers of migrated mouse GPR43KO CD4^+^ T cells treated with 5 mM acetate in an *in vitro* trans-well assay, counted by flow cytometry. Data shown from one of two independent experiments (A-C, E, H-J) and combined three independent experiments (D), one of three independent experiments (F, G), and expressed as mean ± SD. Statistical difference was determined by Student’s two-tailed t-test. *p < 0.05; **p < 0.01; ***p < 0.001; ****p<0.0001.

### Exposure to acetate reduces T cell migration capacity

We next performed a trans-well migration assay of either mouse CD4^+^ or human Jurkat T cells in the presence and absence of acetate. Here we found a significant decrease in trans-well migration of mouse and human derived CD4^+^ T cells (Figure 1F, 1G). To study the potential impact of decreased T cell migration *in vivo*, primary mouse CD4^+^ T cells were pre-treated for 4 hours with 5 mM acetate *in vitro*, loaded with CellTrace^®^ dye and combined at 1:1 ratio of control to acetate treated CD4^+^ T cells, to a total of 6×10^7^ cells. We, then, adoptively transferred 6×10^7^ cells to recipient C57BL/6 wildtype naïve mice not supplemented with alcohol. Two hours post adoptive transfer, we observed about 50% decrease in migration of acetate treated CD4^+^ T cells to spleens of recipient mice compared to control CD4^+^ T cells (Figure 1H). The same observation was true at 48 hours post adoptive transfer (Figure 1I). Next, we asked whether acetate exerts its effects on T cells via G-protein coupled receptor 43 (GPR43), a membrane receptor bound and activated by short-chain fatty acids (SCFA) ^25^. We found GPR43 knock-out (GPR43KO) CD4^+^ T cells also migrate less in an *in vitro* trans-well assay in the presence of acetate (Figure 1J). These data indicate that the exposure to acetate reduces T cell migration capacity *in vitro* and *in vivo*.

### T cells exposed to acetate exhibit reduced total filamentous actin

T cells require a dynamic cytoskeleton: rapid modifications of filamentous actin (F-Actin), for migration, immune synapse formation and activation ^3^. Upon quantification of total F-Actin levels by Alexa-Fluor-488™-conjugated phalloidin staining and flow cytometry analysis in mouse naïve CD4^+^ T cells exposed to 5 mM acetate, we found a dose dependent decrease of F-Actin median fluorescence intensity (MFI) (Figure 2A). Most of the F-Actin is concentrated to the edges of the cell, also referred to as cortical actin or cortical F-actin, where it serves to help cells maintain and modify shape ^26^. Hence, F-Actin deficiency also manifests itself by increased cell deformability measured by real-time deformability cytometry (RT-DC) ^27^. We performed RT-DC and found an increase in T cell deformability upon acetate exposure (Figure 2B). Plotting T cell deformability against cell size sustained the decrease in deformability upon acetate exposure (Figures 2C, 2D). Together these data reveal a reduction of total F-Actin in acetate-exposed T cells.

**Figure 2.**
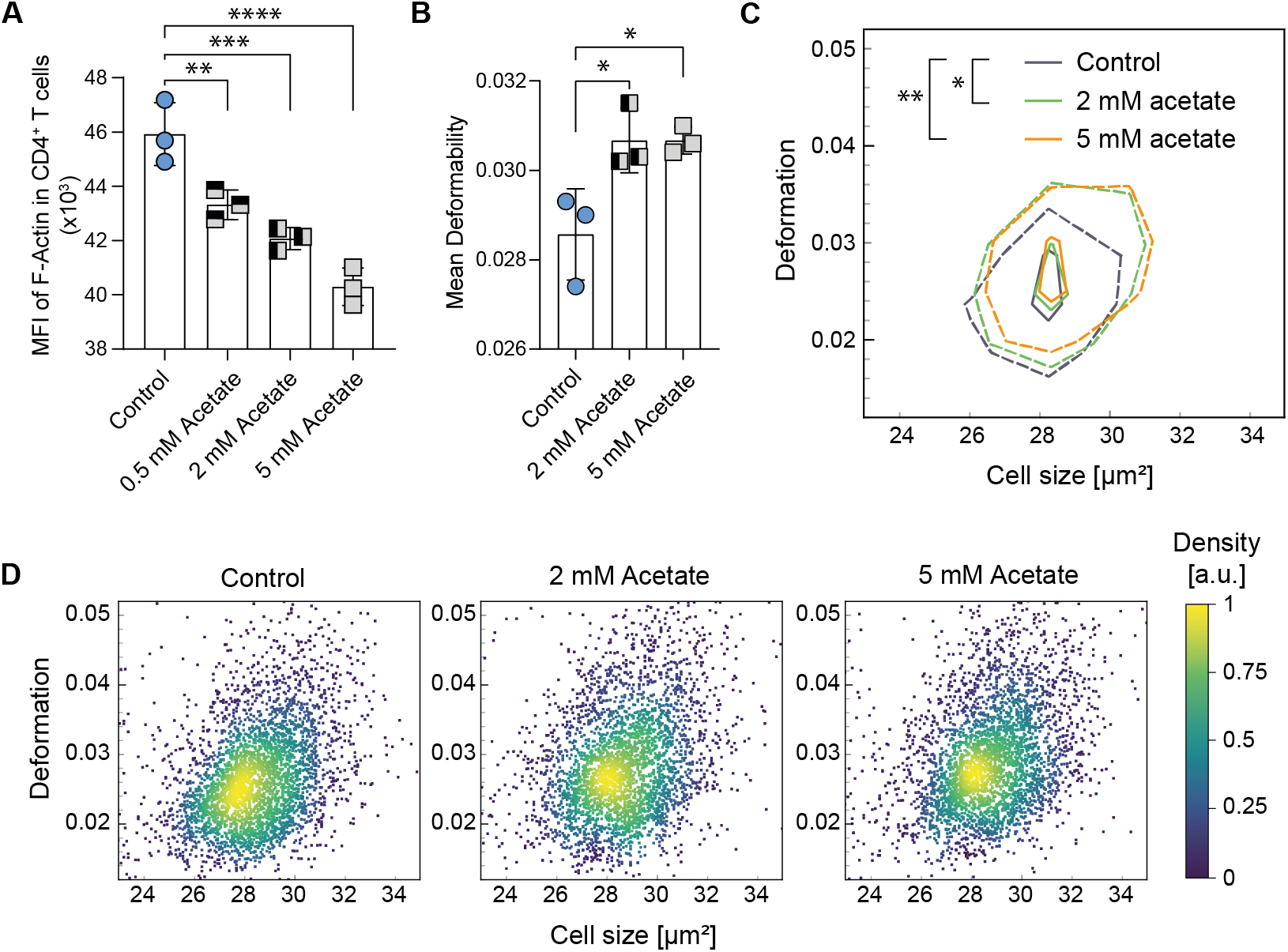
Acetate exposure reduces F-Actin and affects T cell deformability. (**A)** Flow cytometry analysis of mouse CD4^+^ T cells shows a dose-dependent decrease in phalloidin-Alexa Fluor™-488 median fluorescence intensity (MFI). (**B**) A real-time deformability cytometry (RT-DC) of mouse CD4^+^ T cells shows an increase in deformability upon 2 mM acetate and 5 mM acetate treatment (n = 3, CD4^+^ T cells isolated from separate mice). (**C**) Overlay of control, 2 mM and 5 mM acetate treated CD4^+^ T cell deformation and cell size. (**D**) Scatter plots of RT-DC analysis of control and acetate treated (2 mM and 5 mM) naïve CD4^+^ T cell deformation shown in Fig. 2C. Data shown from one of three independent experiments (A) and one of two independent experiments (B, C, D) and expressed as mean ± SD. Statistical difference was determined by one-way ANOVA (A, B). Statistical analyses of panel C were carried out using a one-dimensional linear mixed model that incorporates fixed effect parameters and random effects to analyze differences between cell subsets and replicate variances, respectively. *p*-values were determined by a likelihood ratio test, comparing the full model with a model lacking the fixed effect term. *p < 0.05; **p < 0.01; ***p < 0.001; ****p<0.0001.

### Exposure to acetate increases acetylation of cortactin in T cells

A recent review article by Mu et al. summarized the acetylation of various components of the cytoskeleton ^9^. We identified cortactin, a facilitator and stabilizer of F-Actin branching, as a likely candidate for further research as it has acetylation site in each of its 6.5 repeat regions within the actin-filament binding domain (ABD) ^9^. Initially, it was believed that T cells express hematopoietic homolog of cortactin, hematopoietic lineage cell-specific protein 1 (HS1) ^28^. But later, cortactin was discovered in dendritic cells, macrophages, and lymphocytes ^29^. HS1, in comparison to cortactin, has 3.5 cortactin repeat regions, and acetylation sites on only 2 of those repeats ^30^. In addition, cortactin is a direct target of HDAC6 ^9,31^. As such, treatment of C57BL/6 mice with HDAC6 inhibitor reduced disease severity in a SLE preclinical model and akin to our findings with acetate-exposed collagen induced arthritis (CIA) and T cell dependent vaccination mouse models ^13,14,17^. Cortactin acetylation leads to decreased binding to F-Actin and subsequent F-Actin branching ^31^. We first confirmed cortactin expression in mouse primary CD4^+^ T cells by RNA sequencing (Supplementary Figure 1A). Acetate treatment of Jurkat T cells increased acetylation levels of cortactin as shown by western blotting (Figure 3A). We then identified cortactin protein highly confident (protein score -10lgP-value 189.32) in Jurkat T cells by Mass Spectrometry (ESI MS/MS) and confirmed its acetylation, e.g., at the c-terminal lysine 181 of peptide VDKSAVGFDYQGK (score -10logP 28.34). Because acetylation of cortactin prevents its binding to F-Actin, we mutated lysine residues to arginine residues at the acetylation sites (CTTN_KR) within cortactin ABD ^31^. Microscopy analysis of F-Actin MFI in Jurkat T cells overexpressing CTTN_KR exhibited increased F-Actin amounts even in the presence of acetate (Figure 3B). Further analysis of F-Actin-bound cortactin by Förster Resonance Energy Transfer (FRET) revealed reduced F-Actin-bound cortactin levels upon acetate treatment, and that Jurkat T cells expressing mutated cortactin (CTTN_KR) were resistant to such reduction (Figure 3C). Further, we also quantified acetate’s effect on cortactin-F-Actin binding in HEK293T cells and found a similar result (Figure 3D). Next, in trans-well migration assays we found that the expression of acetylation resistant cortactin (CTTN_KR) rescued acetate induced migration deficiency (Figure 3E). Modifications to F-Actin are required for lamellipodia formation and proper T cell migration ^3^. Cortactin has been shown to be required for lamellipodial persistence and cell migration ^32^. For this reason, we performed 2D Jurkat T cell tracking in cell culture treated wells by live-cell microscopy (Figure 3F). We found that acetate treated Jurkat T cells had a reduced linearity of forward progression and directional change rate, and that cells expressing acetylation resistant cortactin (CTTN_KR) were resistant to the effects of acetate (Figures 3G, 3H). In order to study the highly dynamic lamellipodia formation and persistence, we generated Jurkat T cells expressing LifeAct-mScarlet-i_N1, a short peptide that binds F-Actin. Here too, we observed reduced lamellipodia formation upon acetate treatment of Jurkat T cells (Figure 3I, 3J and Movie S1). Finally, Western Blot analysis of the splenocytes isolated from alcohol-fed mice exhibited increased levels of acetylated cortactin in comparison to water-fed control mice (Figure 3K). In a proof-of-concept experiment, primary cortactin knock-out mouse CD4^+^ T cells exhibited severe impairment of migration in an *in vitro* trans-well assay, exhibiting little known importance of cortactin for proper T cell migration (Figure 3L). These findings indicate that alcohol consumption can directly increase acetylation of cortactin in secondary lymphoid organs such as spleen, and such an increase in T cells impairs T cell migration capacity.

**Figure 3.**
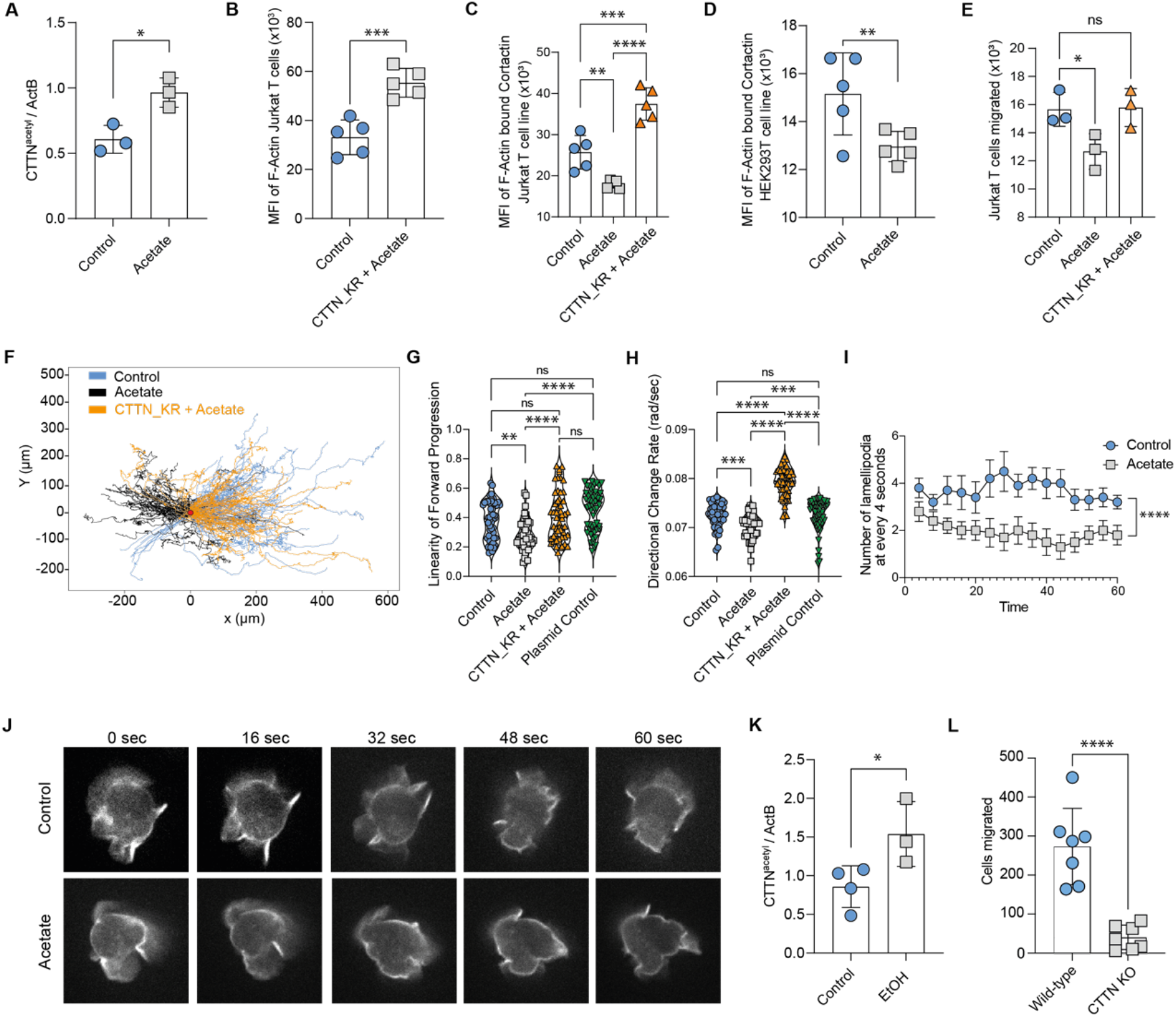
Acetylation resistant cortactin rescues acetate-induced migration deficiency. **(A**) Quantification of western blot analysis of the level of acetylated cortactin in Jurkat T cells treated with 5 mM acetate. (**B**) MFI of F-Actin in acetate treated Jurkat T cells overexpressing CTTN_KR in comparison to control Jurkat T cells. (**C**) MFI of Förster Resonance Energy Transfer (FRET) from F-Actin bound cortactin in Jurkat T cells quantified by confocal scanning laser microscopy in control and acetate treatment, as well as in acetate treated Jurkat T cells expressing acetylation resistant cortactin (CTTN_KR). (**D**) MFI of Förster Resonance Energy Transfer (FRET) from F-Actin bound cortactin in control and acetate treated HEK293T cells quantified by confocal laser scanning microscopy. (**E**) Numbers of migrated Jurkat T cells *in vitro* trans-well assay of control, acetate treated, as well as acetate treated Jurkat T cells expressing acetylation resistant cortactin (CTTN_KR). (**F**) Individual tracks analyzed in panel G and H. (**G**) Linearity of forward progression and (**H**) directional change rate of Jurkat T cells treated with 5 mM acetate, medium control, plasmid electroporation control and Jurkat T cells expressing CTTN_KR and individual cells tracked by Image J Fiji TrackMate plugin ^28, 29^ (n = 50 cells). (**I**) Numbers of lamellipodia per 4-second time points in LifeAct-mScarlet expressing Jurkat T cells upon control and acetate treatment (n = 10 cells). (**J**) Representative still images of control and acetate treated LifeAct-mScarlet expressing Jurkat T cells at 0, 16, 32, 48, and 60^th^ seconds (for full movie see Movie S1). (**K**) Quantification of western blot analysis of the level of acetylated cortactin in the splenocytes of alcohol-fed mice compared to water-fed control (n = 4 control mice, n = 3 alcohol-fed mice (EtOH)). (**L**) Migrated cell count in an *in vitro* trans-well migration assay of primary cortactin knock-out (CTTN KO) mouse CD4^+^ T cells compared to wild-type CD4^+^ T cells, counted by flow cytometry. Data shown from one of three independent experiments (C, E) and one of two independent experiments (A, B, D, F, G, H, K, I, J, L) and expressed as mean ± SD, except for panels G and H where data are expressed as mean ± SEM. Statistical difference was determined by Student’s two-tailed t-test (A, B, D, K, L), one-way ANOVA (C, E, G, H) and area under the curve analysis (I). *p < 0.05; **p < 0.01; ***p < 0.001; ****p<0.0001.

### Acetate-induced F-Actin deficiency impairs T cell activation and proliferation

F-Actin cytoskeleton is required for T cell – antigen presenting cell (APC) interaction and immune synapse formation ^33–35^. Therefore, we performed co-culturing experiment of NP-CGG antigen-pulsed dendritic cells (DCs) and CD4^+^ T cells in order to interrogate if acetate-directed F-Actin deficiency affects T cell activation. T cells were either pre-treated with 5 mM acetate or vehicle, while antigen-pulse of DCs with NP-CGG was performed in an absence of acetate. Later upon combining DCs and T cells, acetate treatment was continued in an effort to replicate *in vivo* acetate exposure within lymphoid organs. Here, flow cytometry analysis revealed reduced surface CD69 levels on acetate treated CD4^+^ CD25^+^ T cells indicating reduced activation (Figure 4A). In the same experiment, flow cytometry analysis of acetate treated CD4^+^ T cells revealed that Ki67 levels, a proliferation marker, were reduced (Figure 4B). Next, we performed confocal fluorescence microscopy to visualize F-Actin at the immune synapse by phalloidin staining and quantified MFI values. There was a decrease in F-Actin accumulation at the immune synapses within T cells between T cells and DCs (Figure 4C). Activation of naïve CD4^+^ T cells with anti-CD3 and anti-CD28 antibody coated beads in the presence of acetate also resulted in reduced F-Actin amounts, as quantified by Alexa-Fluor-488™-phalloidin MFI (Figure 4D). These experiments document the sensitivity of T cells to cytoskeletal deficiencies for migration, immune synapse formation, and activation, and they shed light on our previously published finding of reduced Tfh-B cell conjugate formation upon acetate exposure *in vivo* and *in vitro* ^17^.

**Figure 4.**
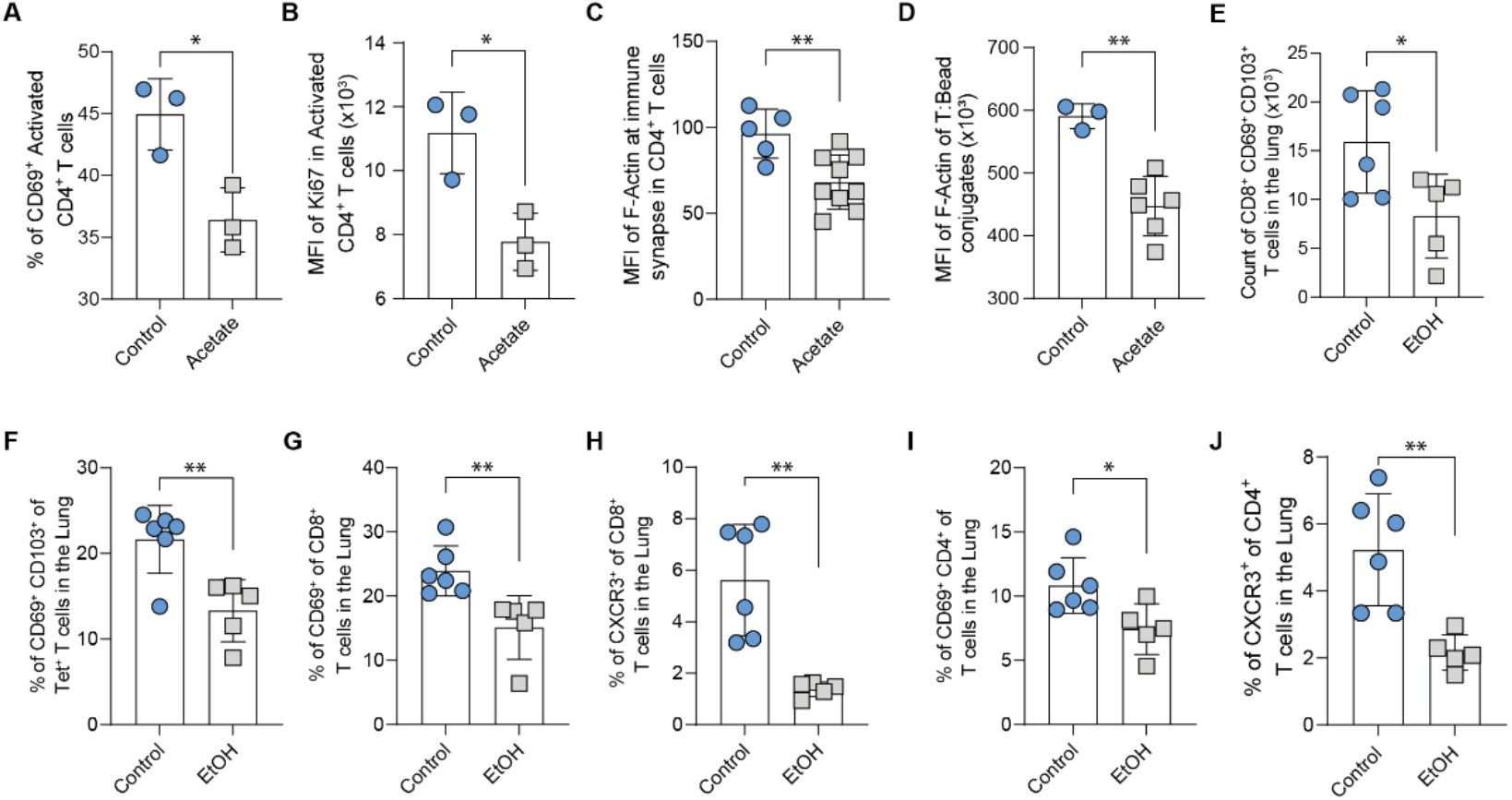
Acetate-induced F-Actin deficiency reduces T cell activation and proliferation. (**A**) Flow cytometry analysis of percentage of CD69^+^ CD25^+^ CD4^+^ and (**B**) Ki67 MFI of CD4^+^ T cells *in vitro* co-culture with NP-CGG pulsed DCs in the presence or absence of 5 mM acetate. (**C**) MFI of F-Actin accumulated in CD4^+^ T cells at the immune synapse with NP-CGG pulsed DCs quantified by confocal laser scanning microscopy and (**D**) with mouse CD4^+^ T cell activator beads quantified by flow cytometry. Flow cytometry analysis of CD69^+^ CD103^+^ (**E**) CD8^+^ T cells and (**F**) influenza specific tetramer (Tet^+^) CD8^+^ T cells in the lungs of alcohol-fed mice in comparison to water-fed control mice. (**G**) Flow cytometry analysis of percentage of CD69^+^ and (**H**) percentage of CXCR3^+^ of CD8^+^ T cells in the lungs of alcohol-fed mice in comparison to water-fed control mice. (**I**) Flow cytometry analysis of percentage of CD69^+^ and (**J**) percentage of CXCR3^+^ of CD4^+^ T cells in the lungs of alcohol-fed mice in comparison to water-fed control mice (each point represents one mouse. n = 6 control, n = 5 alcohol-fed mice). Data shown from one of three independent experiments (A, B, D), two independent experiments (C, E, F, G, H, I, J) and expressed as mean ± SD. Statistical difference was determined by Student’s two-tailed t-test. *p < 0.05; **p < 0.01; ***p < 0.001; ****p < 0.0001.

### Acetate-exposed mice exhibit inhibited T cell responses in the lungs upon influenza infection

Since we have witnessed acetate’s effect on T cell cytoskeleton, we wondered whether alcohol-sourced acetate will impair T cell migration and activation during viral infection. For this, we utilized mouse influenza infection model. Here, alcohol consuming mice were infected with influenza H1N1 and two weeks later lungs isolated and analyzed. Flow cytometry analysis revealed reduced numbers of CD8^+^ CD69^+^ CD103^+^ T cells in the lungs of alcohol consuming influenza infected mice in comparison to water-fed control mice (Figure 4E). Interestingly, we also found decreased percentage of CD69^+^ Tet^+^ influenza specific CD8^+^ T cells and CD69^+^ CD8^+^ T cells in alcohol consuming influenza infected mice in contrast to water-fed control mice (Figures 4F, 4G). In addition, we observed reduced percentage of CXCR3^+^ CD8^+^ T cells in the lungs of alcohol consuming mice in comparison with water-fed control mice (Figure 4H). Flow cytometry analysis of the CD4^+^ T cells in the lungs, revealed a reduction in percentage of CD69^+^ and also CXCR3^+^ CD4^+^ T cells in alcohol consuming mice in contrast to water-fed control mice (Figures 4I, 4J). Together these data reflect deficiencies in T cell migration, activation, and function.

## Discussion

Our findings indicate that alcohol-sourced acetate accumulates in lymphoid organs, increases cellular acetylated cortactin levels preventing F-Actin branching and impeding T cell migration (overview Fig. 5). The Arp2/3 complex is a part of F-Actin branching nucleator complex ^35^. But it does require cortactin to facilitate its binding to F-Actin. Whereas cortactin was shown to increase the affinity of Arp2/3 complex for F-Actin by almost 20-fold ^36^.

**Figure 5.**
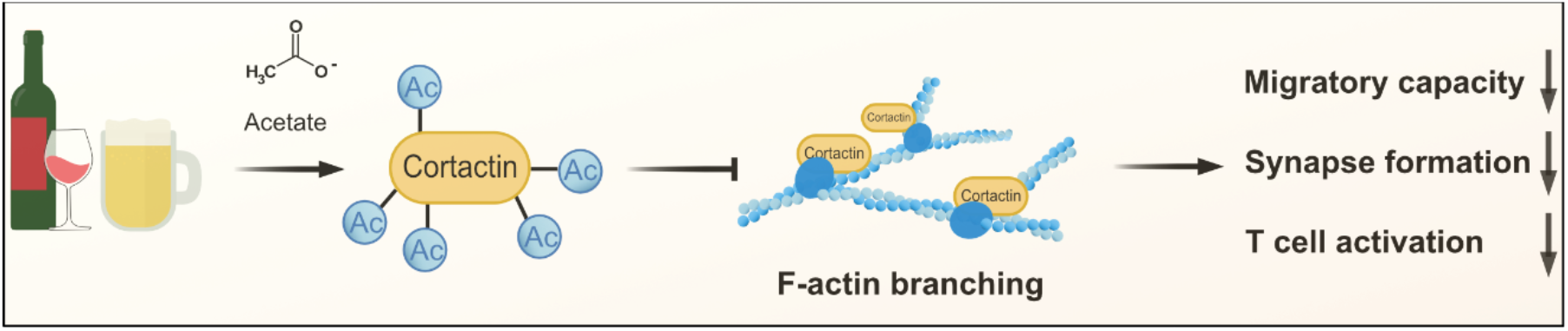
Overview of the mode of acetate’s action on T cell F-Actin reorganization. Alcohol consumption results in accumulation of acetate in lymphoid organs, increases acetylated cortactin levels in T cells. This results in reduced F-Actin branching and consequently reduced migratory capacity, synapse formation, and activation.

In addition, cortactin plays a key role in linking cortical F-Actin to the cell membrane ^37^. Inhibition of Arp2/3 complex formation reduced T cell deceleration upon encountering high affinity antigens to a level observed for low affinity antigens ^38^. This finding potentially explains the weak immune synapse formation in our current study and also reduced T_FH_-B cell conjugate formation upon acetate exposure in our previously published findings ^17^. F-Actin rearrangement enables cells to migrate, direct adhesion molecules to cell-cell contact zones, divide, and regulate various other cellular processes ^39,40^. Cytoskeleton is one of the central elements for optimum T cell function ^41^. For example, functional actin cytoskeleton has been shown to be important for T cell sampling of antigen-MHCII complexes on APCs with subsequent role in immune synapse formation ^33^. CD4^+^ and CD8^+^ T cell migration patterns through secondary lymphoid organs (SLOs) differ, CD4^+^ T cells spend more time probing MHCII molecules presented on DCs and are significantly faster in entering and exiting SLOs than CD8^+^ T cells ^42–44^. It is possible that diminished F-Actin branching can therefore mitigate TCR coupling with high-affinity antigens and reduce T cell arrest and subsequent activation. Interestingly, in a mouse model of type 1 diabetes, mice fed with acetate-yielding chow exhibited a reduction in autoreactive T cells (both CD4^+^ and CD8^+^) and ultimately reduced the incidence of the disease ^45^. In another study of experimental autoimmune encephalomyelitis (EAE), cortactin knock-out mice exhibited decreased incidence, disease severity, and infiltration of the central nervous system by CD4^+^ T cells ^46^. Interestingly, we observed the same effect in alcohol-fed EAE mice ^17^. Similarly, in an influenza infection model, alcohol-fed mice exhibited decreased homing to lungs, reduced CD69, CD103, and CXCR3 positive tissue resident CD8^+^ T cells. Cortactin is overexpressed in B cells of chronic lymphoblastic leukemia patients (CLL), implicated in T cell acute lymphoblastic leukemia (T-ALL), and promotes migration in cancer ^47,48^. Identification of an Achille’s heal of T cell migration, motility, and immune synapse formation can set the stage for pharmacological and/or natural targeting in cases of autoimmunity. Further, such interventions can also help improve immune responses to vaccinations. In an evolutionary perspective, hominids adapted to alcohol metabolism by feeding on fermenting fruits from the forest floor ^49^. It is possible that indirect alcohol consumption mitigated adverse immune responses in early humans under high microbial load. Today, we see the same effects reflected in mitigation of autoimmunity and susceptibility to infections amongst alcohol consumers ^50^.

## Limitations of the study

Although data shown herein directly link alcohol consumption to increased acetylation of cortactin, it is important to note that alcohol-sourced acetate can have other effects on T cell biology. While our study provides a robust result on the new role of cortactin in T cells, there are currently no mouse strains where cortactin is knocked out specifically in T cells, hence our limited use of in vivo models.

## Acknowledgments

We thank Schimpf M., Danzer H. and Steffen F. for technical assistance. We also thank Schmidt K. and Harrer T. for their help with T cell electroporation. We are grateful to Dürholz K., Schälter F., Saad M.S.A., Vogg M., Ramming A., Weidner D. for their support and help with experiments. We thank Geisberger S. and Kempa S. for ^13^C Citrate quantification and data analysis. We thank Tripal P. Winter Z. and Schmidt B. from the OICE in Erlangen for technical expertise. We thank Mielenz D., Sarter K., and Brandl C., for helpful discussions. We thank Prof. Dr. Klemens Rottner from Helmholtz Centre for Infection Research (Braunschweig, Germany) for providing Cortactin Knock-Out mice.

## Author Contributions

Conceptualization: VA, GS, MMZ. Methodology: VA, MF, MH, JH, MK, DL. Investigation: VA, MK, MT, MMZ. Visualization: VA, MF, MK. Funding acquisition: GS, MMZ. Project administration: VA, MMZ. Supervision: MMZ. Writing – original draft: VA, MMZ. Writing – review & editing: VA, MT, JG, GS, MMZ

## Declaration of Interests

Authors declare that they have no competing interests.

**Supplementary Figure 1.**
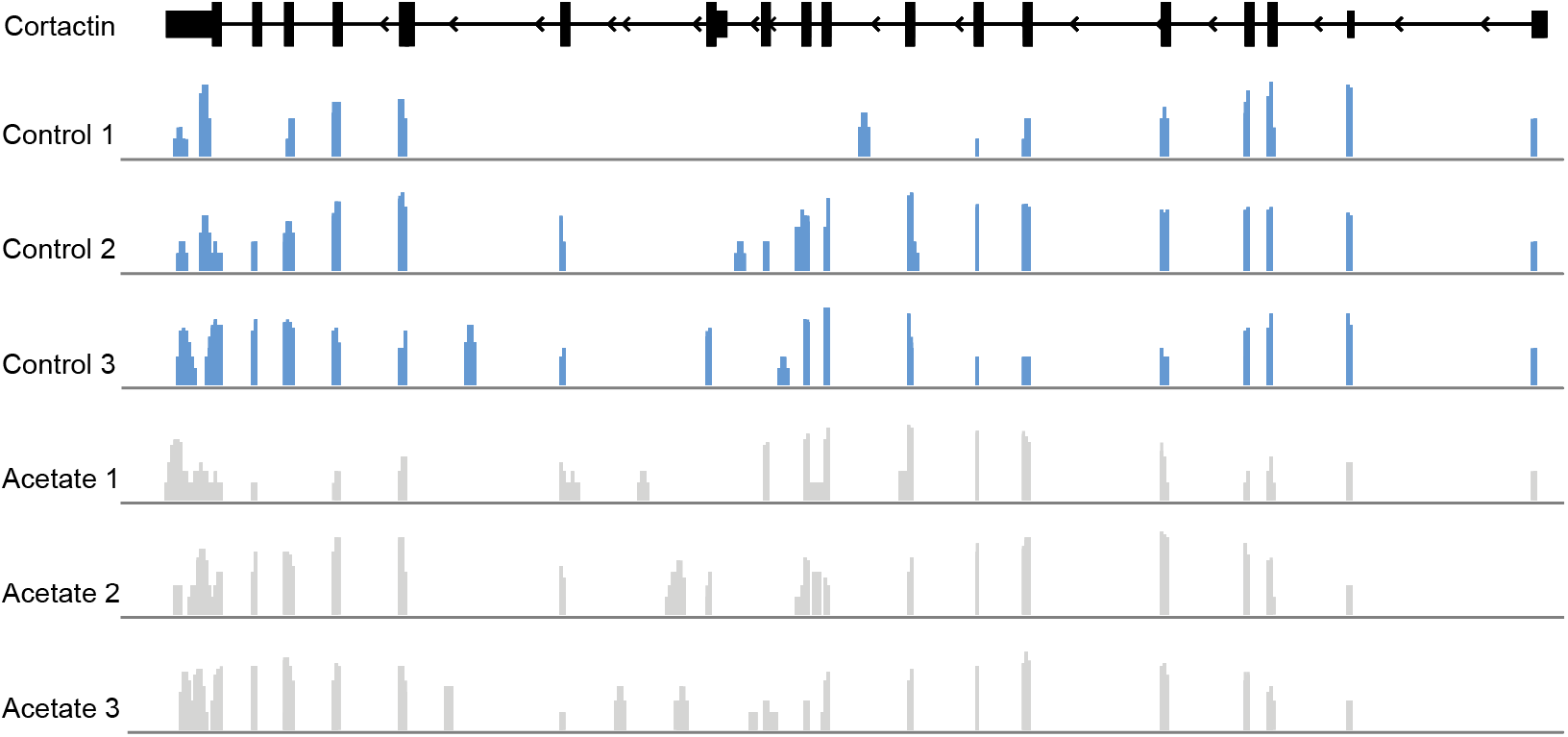
Cortactin is expressed in primary mouse CD4^+^ T cells. Read peaks from RNA-sequencing in the cortactin gene in control (blue) and acetate-treated (grey) T cells are shown.

## Figure Legends

**Movie S1. Jurkat T cells exposed to acetate exhibit reduced lamellipodia formation**.

Jurkat T cells expressing LifeAct-mScarlet treated with 5 mM acetate or vehicle (R10 medium) in R10 medium (n = 10 cells per treatment), visualized at 6 frames per second (each frame = 4 seconds) for 60 seconds. Stills included as Fig. 3J.

## Materials and Methods

### Mice

Female, 6 weeks old C57BL/6NCrl mice were purchased from Charles River (Germany). Mice were co-housed for 3 weeks prior to start of experiments. Cortactin knock out mice were kindly provided by Prof. Dr. Klemens Rottner from Helmholtz Centre for Infection Research (Braunschweig, Germany). All mice were housed, and experiments were conducted under specific pathogen-free conditions. All of the protocols for animal experiments were approved by the local ethics authorities of the Regierung von Unterfranken (55.2-2532-2-703, 55.2.2-2532-2-1515).

### Adoptive transfer experiments

CD4^+^ T cells (naïve or total) were enriched with naïve or total CD4^+^ T cell negative enrichment kits (Stemcell, Biolegend) from the spleens of female, 8 weeks old C57BL/6NCrl mice. T cells were then either treated with 5 mM sodium acetate or sodium matched R10 medium (RPMI 1640, 10% v/v Fetal Bovine Serum, supplemented with 50 μM β-mercaptoethanol, non-essential amino acids, 2 mM L-Glutamine, sodium pyruvate, 10 mM HEPES, and 1% penicillin and streptomycin) for 4 hours. Cells were spun down (300xg) and washed two times with PBS at room temperature (RT). Cells were then loaded with CellTrace® cell dye (CFSE and Violet, for control and acetate treatment) according to manufacturer’s instructions (Thermo Fisher) and incubated further 30 minutes with corresponding treatment (control or acetate containing R10 medium). Cells were then counted and adjusted to 3×10^7^ cells for each treatment and combined to make 6×10^7^ cells in PBS prior to transfer to recipient mice (C57BL/6NCrl, female, matching age) intravenously in the tail vain. Recipient mice were sacrificed 2 hours and 48 hours post adoptive transfer, and spleens collected for flow cytometry.

### NP-CGG Immunizations

Female, 8 weeks old C57BL/6NCrl mice were co-housed and 1 week before immunization, mice were given either water, 10% (v/v) Ethanol (Roche) and 2% (w/v) Glucose (Sigma), all feedings were changed every 3 days. For primary NP-CGG immunization, mice were injected intraperitoneally (i.p.) with 100 µg of NP-CGG (LGC Biosearch Technologies) in 200 µL of alum (Thermo Scientific) according to manufacturer’s instructions. 14 days later, mice were boosted with 100 µg of NP-CGG in 200 µL of alum. One week later, mice were sacrificed, and spleens collected for isolation of CD4^+^ T cells and dendritic cells (Stemcell, Biolegend) for *in vitro* experiments.

### Influenza infection model

Starting 1 week before infection, C57BL/6NCrl mice were given either 2% (w/v) Glucose water or 10% (v/v) Ethanol (Roche) and 2% (w/v) Glucose (Sigma). All feedings were changed every 3 days and were continued throughout the infection. Animals were infected by intranasal instillation of 200 PFU of H1N1 A/Puerto Rico/8/1934 in 50 µL PBS under general anesthesia. Lungs were harvested 14 days post infection. For T cell analyses, lungs were cut in pieces and were treated with 500 units Collagenase D and 160 units DNase I in 2 mL R10 medium (RPMI 1640 supplemented with 10% FCS, 2 mM L-Glutamine, 10 mM HEPES, 50 μM β-mercaptoethanol and 1% penicillin/streptomycin) for 45 minutes at 37°C. Lungs were then mashed through a 70 µm cell strainer before subjected to erythrocyte lysis. One fifth of the cell suspension was stained with a pentamer for NP366-374-specific CD8^+^ T cells (1:40; ProImmune), anti-CD8-BV711 (1:300, clone 53-6.7, Biolegend), anti-CD4-BV605 (1:300, clone RM4-5, Biolegend), anti-CD45-BV510 (1:300, clone 30F11, Biolegend), anti-CD69-PerCP-Cy5.5 (1:200, clone H12F3, Biolegend), anti-CD103-PE (1:200, clone 2E7, Invitrogen), anti-CXCR3-APC-Fire750 (1:100, clone CXCR3-173, Biolegend), anti-CD127-FITC (1:300, clone A7R34, Biolegend), and anti-KLRG1-PE-Cy7 (1:300, clone 2F1, Invitrogen). Data were acquired on an AttuneNxt (ThermoFisher) and analysed using FlowJo™ software (Tree Star Inc.).

### Flow cytometry

Spleens were smashed and filtered through 40 μm gauze (BD Biosciences). Single-cell suspensions were incubated with antibodies for 30 minutes at 4°C, washed and analyzed with Cytoflex S flow cytometer (Beckman Coulter). Flow cytometry analysis of *in vitro* experiments were performed by centrifuging cells in 96-well V-bottom plates at 300xg and 4°C for 5 minutes. Antibody staining for surface markers were performed at 4°C for 30 minutes, followed by two times wash with FACS Buffer (PBS, 2 mM EDTA, 2% Fetal Bovine Serum) and one-time wash with PBS. Cells were then resuspended in 100 µL of PBS and fixed with addition of 100 µL of IC Fixation Buffer (Thermo Fisher) and incubated at 4°C for 15 minutes protected from light. Cells were then washed three times with 1x Permeabilization buffer and incubated for 1 hour at RT for permeabilization (Thermo Fisher) followed by intracellular staining. Intracellular staining of filamentous actin was performed by incubating cells with phalloidin-Alexa-Fluor®-488 (Abcam) at 1:1000 dilution for 2 hours at RT in PBS with 1% w/v BSA, protected from light.

### *In vitro* migration

Jurkat T cells or naïve CD4^+^ T cells were treated with 5 mM sodium acetate containing R10 medium for 4 hours prior to Transwell migration assay. T cells were then seeded at 10^5^ cells in 100 µL total volume above 3 µm and 8 µm pore size transwell inserts (GreinerBio), mouse CD4^+^ T cells and Jurkat T cells, respectively. Transwell inserts were placed in 24-well plate containing R10 medium matching the volume height within the insert (600 µL) and incubated for 4 hours. Migrated cells were collected from the bottom wells and centrifuged, resuspended in 100 µl FACS buffer, quantified by flow cytometry.

### *In vitro* co-culture of Antigen Presenting Cells (APCs) and T cells

Dendritic cells (DCs) and CD4^+^ T cells were enriched from C57BL/6CNrl mice immunized two times, 14 days apart, with 100 µg NP-CGG in 200 µL of Imject™ Alum and sacrificed on day 21 (Stemcell, Thermo Fisher, LGC). CD4^+^ T cells were pre-treated with 0.5, 2.0, or 5 mM sodium acetate or vehicle (R10 medium) containing R10 medium for 4 hours. Meanwhile, DCs were pulsed with 20 µg/mL NP-CGG (LGC) and 100 ng/mL LPS (Sigma Aldrich) in R10 medium for 4 hours. CD4^+^ T cells were adjusted to 10^5^ cells per well of 96-well flat bottom cell culture plate. DCs were then washed two times with fresh medium and combined with CD4^+^ T cells for 48 hours. Final total volume per well was 200 µL. At the conclusion of 48-hour incubation, cells were moved to a 96-well V-bottom cell culture plate, wells flushed and collected. Cells were fixed and stained as explained under Flow Cytometry section.

### *In vitro* activation of CD4^+^ T cells

Naïve CD4^+^ T cells were enriched from C57BL/6CNrl mice (Stemcell). Then, the cells were pre-treated with 0.5, 2.0, or 5 mM sodium acetate or vehicle (R10 medium) containing R10 medium for 4 hours. Dynabeads™ Mouse T-Activator CD3/CD28 beads were added according to manufacturer’s instructions for 4 hours (Thermo Fisher). At the conclusion of 4 hours, cells were centrifuged at 4°C at 200xg for 10 minutes and fixed with IC fixation buffer (Thermo Fisher) in 200 µL (100 µL IC Fixation buffer + 100 µL PBS) for 15 minutes at 4°C. Following fixation, cells were washed with PBS three times and permeabilized with 1x eBiosciences™ permeabilization buffer for 1 hour at room temperature (Thermo Fisher). Phalloidin-iFluor 488 (Abcam) stock solution was diluted in PBS containing 1% w/v BSA 1000x. After permeabilization, cells were washed three times and Phalloidin-iFluor 488 solution added for 2 hours at RT.

### RNA sequencing

Naïve CD4^+^ T cells were treated with 2.0 mM sodium acetate or vehicle (R10 medium) containing R10 medium for 4 hours. RNA was isolated using the RNeasy micro kit (Qiagen). Sequencing was performed on the Illumina platform (Novogene, Europe). Raw reads were processed through fastp (Galaxy). Mapping the processed data to the reference genome (GRCm39) was performed using Star software, and alignment visualization was done with the integrative genomics viewer (IGV).

### Immune synapse formation

Dendritic cells (DCs) and CD4^+^ T cells were enriched from C57BL/6CNrl mice immunized two times, 14 days apart, with 100 µg NP-CGG in 200 µL of Imject™ Alum and sacrificed on day 21 (Stemcell, Thermo Fisher, LGC). CD4^+^ T cells were pre-treated with 0.5, 2.0, 5.0 mM acetate or vehicle (R10 medium) for 6 hours in R10 medium. Simultaneously, DCs were pulsed with 20 µg/mL NP-CGG in R10 medium for 6 hours. Then, 10^6^ pre-treated naïve CD4^+^ T cells and 10^6^ NP-CGG pulsed DCs were combined in 100 µL of R10 medium for 15 minutes at 37°C. Then 1.5 mL of 1.5% paraformaldehyde (PFA) was added to arrest T-DC conjugates. Cells were then washed with PBS+2% FCS (v/v) three times. Cells were then stained for flow cytometry with anti-mouse CD4 BV421™ and anti-mouse CD11c BV510™, or for microscopy anti-CD4 Alexa Fluor 647™ only in PBS+2% FCS (v/v) for 30 minutes at 4°C. Cells were washed, and fixed with eBiosciences™ IC fixation buffer for 15 minutes at 4°C. Cells were washed with PBS three times and permeabilized with 1x eBiosciences™ permeabilization buffer for 1 hour at room temperature. Phalloidin-iFluor 488 stock solution was diluted in PBS containing 1% w/v BSA 1000x. After permeabilization, cells were washed three times and Phalloidin-iFluor 488 solution added for 2 hours at RT. Then, cells were washed with PBS three times with 5-minute incubations in PBS to remove all unbound Phalloidin-iFluor 488. For microscopy samples, nuclei were staining with DAPI in PBS for 5 minutes and then washed and resuspended in 60 µL of PBS. Drops of 30 µL were placed on microscope slides and allowed to dry in the dark. After, ProLong™ Glass Anti-fade mounting medium was used to mount slides and let cure for 24 hours. Samples for flow cytometry were resuspended after washing Phalloidin-iFluor 488 out and resuspending in PBS.

### Live cell tracking

Jurkat T cells and Jurkat T cells overexpressing acetylation resistant cortactin (CTTN_KR) were cultured in R10 medium supplemented with 5 mM sodium acetate for 4 hours. For acetate treatment control, Jurkat T cells were cultured in R10 medium. As a control for plasmid electroporation, Jurkat T cells electroporated with mScarlet_H plasmid were used. The cells were then transferred to incubated and CO2 gassed Zeiss Cell Discoverer (Carl Zeiss) live imaging microscope. Then images were taken by brightfield illumination with Apochromat 10x objective in 2-minute intervals for a total of 180 minutes. Cells were then tracked using TrackMate plugin for Fiji ^51,52^. To visualize individual tracks, tracks were obtained from TrackMate and re-centered at the origin of the plot.

### Live cell imaging of lamellipodia

Jurkat T cells overexpressing pLifeAct_mScarlet-i_N1 (see Electroporation methods), were treated with 5 mM sodium acetate or vehicle (R10 medium) containing R10 medium for 4 hours. Later, cells were seeded into µ-Slide 8 well microscope slides (ibidi) and placed into incubated microscopy chamber. Zeiss Spinning Disc Axio observer Z1 microscope with oil immersed 63x 1.2 NA objective and EVOLVE 512 EMCCD with Yokogawa CSU-X1M 5000 was used to locate cells. Cells were illuminated with a laser at 561 nm wavelength, and emissions captured through band-pass filter 629/62 every 4 seconds for 60 seconds. Images were later processed and analyzed for numbers of lamellipodia at each time point and counted. Statistical analysis was done by area under the curve analysis.

### Cortactin lysine to arginine mutagenesis

Recombinant cortactin gene with lysine residues 87, 124, 161, 198, 235, 272, 309, and 319 replaced with arginine, was synthesized, and cloned into pcDNA3.1plus vector by GeneArt™ Thermo Fisher. Jurkat T cells were electroporated as described below and clones were selected with 800 µg/mL G418 (InvivoGen) antibiotic for 7 passages.

### Electroporation of Jurkat T cells

Jurkat T cells were maintained in R10 medium. 10^7^ Jurkat T cells were washed two times with RPMI1640 medium only, centrifugation steps at 200xg for 10 minutes, at RT. Jurkat T cells were then washed one more time with 5 mL of OptiMEM® medium and resuspended in 100 µL of OptiMEM® medium. 25 µg (1 µg/µL concentration) of plasmid DNA was added to electroporation 4 mm cuvettes (Bio-Rad) first and then cells added to the cuvettes. Electroporation was carried out with Bio-Rad Gene Pulser Xcell Electroporation Systems at Square wave, 500 Volts, 3 ms, Pulse 1, Interval 0). After the electroporation, cells were transferred to T25 cell culture flasks with 5 mL of prewarmed (37°C) R10 medium and incubated overnight. For control purposes, no plasmid DNA containing cell mixture was used for electroporation.

### Real-time Deformability Cytometry (RT-DC)

RT-DC measurements were performed using an AcCellerator instrument (Zellmechanik Dresden). T cells were resuspended in measurement buffer composed of 0.6% (m/v) methyl cellulose dissolved in phosphate-buffered saline adjusted to a viscosity of 60 mPa s at 24°C using a falling ball viscometer (Haake; Thermo Fisher Scientific). The suspension of T cells was loaded into a 1 mL syringe attached to a syringe pump and connected by PEEK-tubing (IDEX Health & Science) to a microfluidic chip made of PDMS bonded on cover glass. A second syringe with pure measurement buffer was attached to the chip. The microfluidic chip consisted of two inlets and one outlet. The measurement was performed in a narrow channel constriction of 15 × 15 μm square cross section. The total flow rate was 0.024 μL/s, with sheath flow rate of 0.018 μL/s and sample flow rate of 0.006 μL/s. Measurement temperature was 28°C. Images were acquired at a frame rate of 1250 fps. Cells were detected in a region of interest of 250 × 80 pixels. The cell images were analyzed using the analysis software Shape-Out version 2.3.0 (available at https://github.com/ZELLMECHANIK-DRESDEN/ShapeOut2) and Python 3.7 using dclab library. We applied a gate for cross-sectional cell size (15-35 μm^2^) and for the area ratio (1– 1.05). The calculation of deformation, a measure of how much the cell shape deviates from circularity, was obtained from the image using the projected area and cell contour length calculated from the convex hull. Statistical analyses were carried out using a one-dimensional linear mixed model that incorporates fixed effect parameters and random effects to analyze differences between cell subsets and replicate variances, respectively. p-values were determined by a likelihood ratio test, comparing the full model with a model lacking the fixed effect term.

### Protein Isolation and Western Blot

After acetate treatment of Jurkat T cells, naïve CD4+ T cells activated either with 0.5 µg/mL PMA, 1 µg/mL ionomycin or NP-CGG-pulsed CD11c+ DCs then depleted off DCs, T cells were washed two times with ice-cold PBS containing 5 mM sodium butyrate, and nuclear/cytoplasmic protein was extracted according to manufacturer’s instructions (Thermo Fisher). Protein fractions were then frozen and stored at -80°C until analyzed by western blotting. Protein extracts were separated on 4-12% Bis-Tris gradient SDS-polyacrylamide gel, then transferred onto PVDF membrane, blocked with 5% milk in TBS 0.05% Tween 20 for 1 hour. Then membranes were probed with anti-cortactin, anti-acetyl cortactin, anti-GAPDH, anti-beta-actin, anti-histone 3, anti-acetyl-lysine-27-histone-3, and anti-histone 3 antibodies, and visualized with appropriate HRP-conjugated secondary antibody. The signal acquired from chemiluminescence (Celvin S) for protein of interests were optimized to loading control and quantified with Image J blot analysis tool.

### Proteomic analysis of Cortactin

For proteomic samples containing about 2 mg of total protein Jurkat T cells were treated either with 5mM acetate in R10 medium or without acetate as the control. Cells were washed three times with ice cold PBS, lysed with sonication, and cleared by Ultracentrifugation. Mass spectrometry (ESI/LC/MS/MS) was performed as described ^53^. In short, peptides for a topdown proteomic approach were prepared applying filter-aided sample preparation (FASP) ^54^. After reduction and alkylation on a 10kDa membrane filter a total of 1.5 µg of protein of each sample was digested with 1µg trypsin for 18h at 37ºC. Resulting peptides were extracted by centrifugation and desalted on C18 stage tips. Prior to MS analysis, tryptic peptides were dried under vacuum and resuspended in 15µl of 10% formic acid (FA). MS analysis was carried out as described ^53^. In short, resulting peptides (approx. 1 µg) were loaded on a nanoflow Ultimate 3000 HPLC (Dionex, Sunnyvale, CA, USA) for separation on an EASY-Spray column (Thermo Fisher Scientific; C18 with 2 μm particle size, 50 cm × 75 μm), with a flow rate of 200 nL/min by increasing acetonitrile concentrations over 120 min. All samples were analyzed on an Orbitrap Fusion (Thermo Fisher Scientific, Waltham, MA, USA) with the previously described MS/MS settings ^53^. ESI spray voltage was 2.0 kV, scan range 300–2000 (*m*/*z*), with a maximum injection time of 50 ms and an AGC target of 400 k for the first stage of mass analysis (MS^1^). The most intense ions were selected for collision-induced dissociation with a collision energy of 35%, a maximum injection time of 250 ms, and an AGC target of 100 for the second stage of mass analysis (MS^2^). All raw files were analyzed using PEAKS Studio 8.5 (Bioinformatics Solutions, Waterloo, Ontario, Canada; ^55^) and searches were performed against the *Homo sapiens* uniprot database downloaded in July 2019. Oxidation of methionine and acetylation of lysin were set as dynamic modifications and carbamidomethylation of cysteines as static modification, Parent mass tolerance was set to 10.0 ppm and fragment mass tolerance to 0.5 Da for tryptic peptides allowing a maximum of two missed cleavages. The MS proteomics data will be deposited to the ProteomeXchange Consortium via the PRIDE ^56^ partner repository upon completion of peer-review.

### Histone extraction

Naïve CD4^+^ T cells were pre-treated with 0.5, 2.0, 5.0 mM sodium acetate or vehicle (R10 medium) containing R10 medium for 4 hours. Cells were then washed with ice cold PBS supplemented with 5 mM sodium butyrate to preserve histone acetylation. Then the cytoplasm was extracted by resuspending cells in ice cold extraction buffer at 10^7^ cell/mL, incubating on ice for 5 minutes, and centrifuging at 6500xg for 10 minutes at 4°C. Supernatant was discarded and the cytoplasm extracted for the second time. Histones were extracted by resuspending the pellet in 0.25 M HCl at a density of 4×10^7^ cells/mL. Samples were then sonicated for 30 seconds, placed on ice, then sonicated for further 30 seconds. Afterwards, tubes were placed on rollers at 4°C for 1 hour. Then, samples were centrifuged at 12000xg for 10 minutes at 4°C. Supernatant was collected and neutralized with 2M NaOH at 1/10th of the volume of the supernatant. Extracted histones were frozen and stored at -80°C until analyzed by western blotting.

### Protein isolation from splenocytes

Spleens of alcohol-fed and water-fed C57BL/6NCrl mice were isolated and placed in ice cold PBS containing 5 mM sodium butyrate after 3 weeks of feeding at steady-state. Spleens were smashed and filtered through 40 μm gauze (BD Biosciences). Single-cell suspensions were spun down and resuspended in 5 mL of 1x Red Blood Cell Lysis buffer (Biolegend) and incubated on ice for 5 minutes with occasional swirl. Then, 25 mL of ice-cold PBS was added, and cells centrifuged at 300xg for 10 minutes. Cells were resuspended in 10 mL ice-cold PBS and spun down. Protein isolation of cytoplasmic and nuclear fraction was performed with NE-PER™ Nuclear Cytoplasmic Extraction (Thermo Fisher) supplemented with Pierce™ Protease inhibitor cocktail tablet (Thermo Fisher). Isolated proteins were kept at -80°C until analyzed by western blotting.

### ^13^C Citrate quantification

Label incorporation into citrate was analyzed by gas chromatography (GC – MS). In brief, cells were subjected to methanol-chloroform-water extraction and polar phase was dried under vacuum. For derivatization, 20 µL per sample of 40 mg methoxamine per mL pyridine were added and incubated at 30°C for 90 minutes, followed by the addition of 80 µL per sample of MSTFA, incubated at 37°C for 60 minutes. GC-MS analysis was performed on an Agilent 7890B GC – Pegasus 4D-C GCxGC TOF MS, coupled to a Gerstel autosampler. Samples were injected in splitless mode (injection volume 1 µL). The following temperature program was applied during sample injection: initial temperature of 80°C for 3 seconds followed by a ramp of 7°C/seconds to 210°C and final hold for 3 minutes. Gas chromatographic separation was performed with a Rxi-5MS column (30 m length, 250 µm inner diameter, 0.25 µm film thickness (Restek). Helium was used as carrier gas with a flow rate of 1.2 mL/minute. Gas chromatographic separation was performed with the following temperature gradient: 2 minutes initial hold at 68°C, first temperature gradient with 5°C/minute up to 120°C, second temperature gradient with 7°C/minute up to 200°C, and a third temperature gradient with 12°C/minute up to 320°C with a final hold for 6 minutes. The spectra were recorded in a mass range of 60 to 600 m/z with an acquisition rate of 20 spectra/second. Data was analyzed with ChromaTOF (Leco) and MetMax software ^57^.

### Short-Chain Fatty Acid (SCFA) measurements

4-5 biological replicates of each treatment (control or ethanol feeding) mouse spleens were placed directly into diethyl ether and weighed in a 2 mL polypropylene tube. The tubes were kept in a cool rack throughout the extraction. 50 µL 33% HCl was added, and samples were vortexed for 1 minute and centrifuged for 3 minutes at 4°C. The organic phase was transferred into a 2 mL gas chromatography (GC) vial. For the calibration curve, 100 μl of SCFA calibration standards (Sigma) were dissolved in water to concentrations of 0, 0.5, 1, 5 and 10 mM and then subjected to the same extraction procedure as the samples. For GCMS analysis 1 μl of the sample (4-5 replicates) was injected with a split ratio of 20:1 on a Famewax, 30 m x 0,25 mm iD, 0.25 μm df capillary column (Restek). The GC-MS system consisted of GCMS QP2010Plus gas chromatograph/ mass spectrometer coupled with an AOC20S autosampler and an AOC20i auto injector (Shimadzu). Injection temperature was 240°C with the interface set at 230°C and the ion source at 200°C. Helium was used as carrier gas with constant flow rate of 1 mL/min. The column temperature program started with 40°C and was ramped to 150°C at a rate of 7 °C/min and then to 230°C at a rate of 9 °C/min, and finally held at 230°C for 9 minutes. The total run time was 40 minutes. SCFA were identified based on the retention time of standard compounds and with the assistance of the NIST 08 mass spectral library. Full scan mass spectra were recorded in the 25-150 m/z range (0.5 s/scan). Quantification was done by integration of the extracted ion chromatogram peaks for the following ion species: m/z 45 for acetate eluted at 7.8 minutes, m/z 74 for propionate eluted at 9.6 minutes, and m/z 60 for butyrate eluted at 11.5 minutes. GCMS solution software was used for data processing.

### Plasmids

pLifeAct_mScarlet-i_N1 (Addgene plasmid # 85056 ; http://n2t.net/addgene:85056; RRID:Addgene_85056) and pmScarlet-H_C1 (Addgene plasmid # 85043; http://n2t.net/addgene:85043 ; RRID:Addgene_85043) was a gift from Dorus Gadella ^58^. CTTN_KR plasmid was synthesized and cloned into pcDNA3.1plus vector by GeneArt™ Thermo Fisher.

## Statistical analysis

Data are expressed as mean±SD unless otherwise indicated in the figure legend. Analysis was performed using a two-sided Student’s t test, single comparison or analysis of variance test for multiple comparisons (one-way or two-way ANOVA followed by Tukey’s or Bonferroni’s multiple comparisons test, respectively). All experiments were conducted at least two times, unless otherwise indicated in the figure legends. n-numbers denote number of individual animals or cells isolated from individual animals. P-values of 0.05 were considered significant and are shown as p < 0.05 (*), p < 0.01 (**), p < 0.001 (***), p < 0.0001 (****). Graph generation and statistical analyses were performed using Prism version 8 software (GraphPad).

## Data availability

The data that support the findings of this study are available from the corresponding author upon reasonable request by email to.

## Materials

### Mice

| Strain | Genotype | Supplier |
| --- | --- | --- |
| C57BL/6NCrl | WT | Charles River, Wilmington, USA |
| C57BL/6J | Cortactin -/- | Prof. Dr. Klemens Rottner,<br>Braunschweig, Germany |

### Cell lines

| Cell line | Species | Supplier |
| --- | --- | --- |
| T cell leukemia JURKAT<br>(#ACC 282) | Human | Leibniz Institute DSMZ German<br>Collection of Microorganisms and Cell<br>Cultures GmbH, Braunschweig, Germany |

### Cell Culture

| Reagent | Supplier |
| --- | --- |
| MEM Alpha Medium (1X) + GlutaMAX<br>(#32571-028) | Gibco™ by Thermo Fisher Scientific,<br>Waltham, USA |
| RPMI-1640 (#21875-091) | Gibco™ by Thermo Fisher Scientific,<br>Waltham, USA |
| Opti-MEM™ I Reduced Serum Medium<br>(#31985070) | Gibco™ by Thermo Fisher Scientific,<br>Waltham, USA |
| Dulbecco's Phosphate Buffered Saline (PBS)<br>(#14190-094) | Gibco™ by Thermo Fisher Scientific,<br>Waltham, USA |
| Fetal Bovine Serum (FBS) (#16000-044) | Gibco™ by Thermo Fisher Scientific,<br>Waltham, USA |
| Penicillin Streptomycin (#15140-122) | Gibco™ by Thermo Fisher Scientific,<br>Waltham, USA |
| MEM non-essential amino acids (#M7145-100 mL) | Merck KGaA Darmstadt, Germany |
| Sodium Pyruvate (100 mM) (#11360088) | Gibco™ by Thermo Fisher Scientific,<br>Waltham, USA |
| 2-Mercaptoethanol (50 mM) (#31350010) | Gibco™ by Thermo Fisher Scientific,<br>Waltham, USA |

### Antibodies

| Antibody | Supplier |
| --- | --- |
| Rat Anti-mouse CD11b, PE/Cy7 (#552850) | BD Biosciences, San Jose,<br>USA |
| Rat Anti-mouse CD19, PE/Cy7 (#115520) | Biolegend, San Diego, USA |

| **Antibody** | **Supplier** |
| --- | --- |
| Armenian Hamster Anti-mouse CD3ε, FITC (#100306) | Biolegend, San Diego, USA |
| Rat Anti-mouse CD4, PerCP/Cy5.5 (#100540) | Biolegend, San Diego, USA |
| Rat Anti-mouse CD4, Alexa Fluor® 647 (#100424) | Biolegend, San Diego, USA |
| Rat Anti-mouse CD3, BV510 (#100234) | Biolegend, San Diego, USA |
| Rat Anti-mouse CD4, FITC (#100406) | Biolegend, San Diego, USA |
| Rat Anti-mouse CD25, PE (#102008) | Biolegend, San Diego, USA |
| Rat Anti-mouse FOXP3, BV421 (#126419) | Biolegend, San Diego, USA |
| Rat Anti-human CD4, FITC (#357405) | Biolegend, San Diego, USA |
| Rat Anti-mouse B220, FITC (#103206) | Biolegend, San Diego, USA |
| Mouse Anti-mouse β-actin, HRP linked (#ab49900) | Abcam, Cambridge, United Kingdom |
| Rat Anti-mouse CD8, FITC (#553031) | BD Biosciences, San Jose, USA |
| Rat Anti-mouse CD62L, APC (#104412) | Biolegend, San Diego, USA |
| Rat Anti-mouse CD44, PE (#103008) | Biolegend, San Diego, USA |
| Rat Anti-mouse CD4, Alexa Fluor 647 (#100424) | Biolegend, San Diego, USA |
| Rat Anti-mouse CD45R/B220, Alexa Fluor 488 (#103225) | Biolegend, San Diego, USA |
| Rabbit Anti-mouse Histone 3, unconjugated (#ab1791) | Abcam, Cambridge, United Kingdom |
| Rabbit Anti-mouse Histone 3 Lysine 27 Acetyl, unconjugated (#ab4729) | Abcam, Cambridge, United Kingdom |
| Donkey Anti-rabbit IgG H&L, (HRP) (#ab6802) | Abcam, Cambridge, United Kingdom |
| Mouse Purified anti-Cortactin (CTTN) Antibody (#868102) | Biolegend, San Diego, USA |
| Rabbit Anti-acetyl Cortactin Antibody (#09-881) | Merck, Darmstadt, Germany |
| Goat anti-mouse IgG Alexa Fluor® 594 (#405326) | Biolegend, San Diego, USA |

### Kits

| **Kit** | **Supplier** |
| --- | --- |
| EasySep Mouse naïve CD4<sup>+</sup> T cell Negative Selection kit | Stemcell Technologies, Vancouver, Canada |
| EasySep Mouse CD11c Positive Selection Kit II | Stemcell Technologies, Vancouver, Canada |
| EasySep Mouse CD4<sup>+</sup> Isolation Kit | Stemcell Technologies, Vancouver, Canada |
| NE-PER™ Nuclear and Cytoplasmic Extraction Reagents (#78833) | Thermo Fisher Scientific, Waltham, USA |
| MojoSort™ Mouse CD4 T cell Isolation Kit (#480033) | Biolegend, San Diego, USA |

### Buffers

| Buffer | Composition |
| --- | --- |
| Phosphate-Buffered Saline (PBS) | 137 mM NaCl<br>2.7 mM KCl<br>10 mM Na<sub>2</sub>HPO<sub>4</sub><br>2 mM KH<sub>2</sub>PO<sub>4</sub><br>pH 7.4 |
| Tris-Buffered Saline (TBS) | 50 mM Tris<br>150 mM NaCl<br>pH 7.4 |
| Western Blotting Running Buffer | 3.5 mM SDS<br>25 mM Tris<br>192 mM glycine |
| Western Blotting Transfer Buffer | 25 mM Tris<br>192 mM glycine<br>20% methanol |
| FACS Buffer | Phosphate-Buffered Saline (PBS)<br>2% FCS<br>1 mM EDTA |
| Cytoplasm Extraction Buffer | 25 mM Tris-HCl<br>1 mM EDTA<br>1% NP-40<br>10 mM Sodium Butyrate |

### Chemicals, proteins, and reagents

| Reagent | Supplier |
| --- | --- |
| Bovine Serum Albumin (BSA) (#A7030) | Sigma-Aldrich, Saint-Louis, USA |
| Diethyl Ether (#346136) | Honeywell Research Chemicals,<br>Seelze, Germany |
| di-Sodium hydrogen phosphate dihydrate (#4984.1) | Carl Roth, Karlsruhe, Germany |
| Ethylenediaminetetraacetic acid (EDTA) (#E9884) | Sigma-Aldrich, Saint-Louis, USA |
| Lipopolysaccharide from Escherichia coli O55:B5 (#L6529-1MG) | Sigma-Aldrich, Saint-Louis, USA |
| Ethanol 70% (#T913.3) | Carl Roth, Karlsruhe, Germany |
| Paraformaldehyde (#104005) | Merck, Darmstadt, Germany |
| peqGOLD Trifast (#30-2010) | VWR International, Radnor, USA |
| Potassium chloride (#6781.3) | Carl Roth, Karlsruhe, Germany |
| Potassium dihydrogen phosphate (#3904.2) | Carl Roth, Karlsruhe, Germany |
| Powdered milk (#T145.3) | Carl Roth, Karlsruhe, Germany |
| 2-Propanol (#9866.6) | Carl Roth, Karlsruhe, Germany |
| Sodium acetate (#X891.2) | Carl Roth, Karlsruhe, Germany |
| Sodium chloride (#3957.1) | Carl Roth, Karlsruhe, Germany |
| Sodium butyrate (#303410) | Sigma-Aldrich, Saint-Louis, USA |
| Sodium propionate (#P5436) | Sigma-Aldrich, Saint-Louis, USA |
| Tris(hydroxymethyl)aminomethane (TRIS) (#4855.2) | Carl Roth, Karlsruhe, Germany |
| Tween 20 (#9127.2) | Carl Roth, Karlsruhe, Germany |
| Imject™ Alum Adjuvant (#77161) | Thermo Fisher Scientific, Waltham, USA |
| NP-CGG (#N-5055E-5) | LGC Biosearch Technologies, Berlin, Germany |
| Ovalbumin 323-339 (#vac-isq) | InvivoGen, Toulouse, France |
| Phalloidin, iFluor 488 (#ab176753) | Abcam, Cambridge, United Kingdom |
| Rotipuran 100% Ethanol Pure (#9065.2) | Carl Roth, Karlsruhe, Germany |
| D (+)-Glucose (#HN06.3) | Carl Roth, Karlsruhe, Germany |
| Cell activation cocktail without Brefeldin A (#423302) | Biolegend, San Diego, USA |
| ProLong™ Glass Antifade Mountant (P36984) | Thermo Fisher Scientific, Waltham, USA |
| Tissue-Tek® O.C.T.™ Compound (#4583) | Sakura Finetek Europe B.V., Alphen aan den Rijn, The Netherlands |
| eBiosciences™ IC Fixation Buffer (#00-8222-49) | Thermo Fisher Scientific, Waltham, USA |
| eBiosciences™ IC Fixation & Permeabilization Buffer Set (#88-8824-00) | Thermo Fisher Scientific, Waltham, USA |
| Pierce™ Protease Inhibitor Tablet (#A32961) | Thermo Fisher Scientific, Waltham, USA |
| SuperSignal™ West Femto Maximum Sensitivity Substrate (#34094) | Thermo Fisher Scientific, Waltham, USA |
| Dynabeads™ Mouse T-Activator CD3/CD28 for T cell Expansion and Activation (#11452D) | Thermo Fisher Scientific, Waltham, USA |
| CellTrace™ Violet Cell Proliferation Kit (#C34557) | Thermo Fisher Scientific, Waltham, USA |
| CellTrace™ CFSE Cell Proliferation Kit (#C34554) | Thermo Fisher Scientific, Waltham, USA |
| eBiosciences™ Permeabilization Buffer 10x (#00-8333-56) | Thermo Fisher Scientific, Waltham, USA |
| eBioscience™ IC Fixation Buffer (#00-8222-49) | Thermo Fisher Scientific, Waltham, USA |
| DAPI, Dihydrochloride - CAS 28718-90-3 – Calbiochem (#268298-10MG) | Merck, Darmstadt, Germany |
| G418, Geneticin, solution (#ant-gn-1) | InvivoGen, Toulouse, France |
| RBC lysis buffer (#420301) | Biolegend, San Diego, USA |

### Consumables

| **Consumable material** | **Supplier** |
| --- | --- |
| 1 mL Inject F (#9166917V) | Braun, Kronberg, Germany |
| Amber Vials 12x32x4.6 mm (#27000) | Supelco by Sigma-Aldrich, Saint-Louis, USA |
| Autoclavable Trash Bags (#E706.1) | Carl Roth, Karlsruhe, Germany |
| Autoclave Tap (#3FKLB410104) | Stericlin, Feuchtwangen, Germany |
| Cell culture Dish, PS, 100/22 MM (#664160) | Greiner Bio-One, Kremsmünster, Austria |
| Cell culture microplate 96 well, PS, F-Bottom (#655180) | Greiner Bio-One, Kremsmünster, Austria |
| Cell culture microplate 96 well, PS, U-Bottom (#650180) | Greiner Bio-One, Kremsmünster, Austria |
| Cell culture microplate 96 well, PS, V-Bottom (#651180) | Greiner Bio-One, Kremsmünster, Austria |
| Cell culture multiwell plate, 24 well, PS (#662160) | Greiner Bio-One, Kremsmünster, Austria |
| Cell culture multiwell plate, 12 well, PS (#665180) | Greiner Bio-One, Kremsmünster, Austria |
| Cell culture multiwell plate, 6 well, PS (#657160) | Greiner Bio-One, Kremsmünster, Austria |
| ThinCert® cell culture insert for 24 well plate, 3 µm pore size (#662630) | Greiner Bio-One, Kremsmünster, Austria |
| ThinCert® cell culture insert for 24 well plate, 8 µm pore size (#662638) | Greiner Bio-One, Kremsmünster, Austria |
| µ-Slide 8 Well ibiTreat (#80826) | ibidi GmbH, Gräfelfing, Germany |
| Cell Scraper 25 cm (#831830) | Sarstedt, Nümbrecht, Germany |
| Cell Strainer 70 µm Nylon (#352350) | Corning, Corning, USA |
| Discardit II 2 mL (#300928) | Becton Dickinson, Franklin Lakes, USA |
| Discardit II 5 mL (#309050) | Becton Dickinson, Franklin Lakes, USA |
| Disposable Scalpel No. 10 (#02.010.30.010) | Feather, Osaka, Japan |
| Falcon 50 mL Polypropylene Conical Tube (#352070) | Corning, Corning, USA |
| Falcon 15 mL Polypropylene Conical Tube (#352196) | Corning, Corning, USA |
| Falcon Cell Strainer, 70 µm Nylon (#352350) | Corning, Corning, USA |
| MaxiSorp 96 well plate (#44-2404-21) | Thermo Fisher Scientific, Waltham, USA |
| Microlane 3 20G x 1 ½ - Nr.1 (#301700) | Becton Dickinson, Franklin Lakes, USA |
| Microlane 3 27G x ¾ - Nr.20 (#302200) | Becton Dickinson, Franklin Lakes, USA |
| Microtainer SST Tubes (#365968) | Becton Dickinson, Franklin Lakes, USA |
| Minisart Syringe Filter, 28 mm, Pore Size 0.2 µm, Sterile (#9649101) | Sartorius, Göttingen, Germany |
| Nitrocellulose membrane (#LC2001) | Thermo Fisher Scientific, Waltham, USA |
| PVDF Transfer Membrane, 0.45 µm, 10x10 cm (#88585) | Thermo Fisher Scientific, Waltham, USA |
| Nunc MaxiSorp Flat-Bottom (#44-2402-21) | Invitrogen, Carlsbad, USA |
| Omifix 40 Duo U-40 Insulin (#9161333V) | Braun, Kronberg, Germany |
| Osteo Assay Plate, 24 well (#3987) | Corning, Corning, USA |
| Parafilm PM-996 (#P7793-1EA) | Sigma-Aldrich, Saint-Louis, USA |
| PCR-Strips 8 tubes (#CH88-1) | Carl Roth, Karlsruhe, Germany |
| Safe-Lock Tube 1.5 mL (#0030120.86) | Eppendorf, Hamburg, Germany |
| Safe-Lock Tube 2 mL (#0030120.094) | Eppendorf, Hamburg, Germany |
| SafeSeal SurPhob Tips 10 µL Sterile (#VT0200) | Biozym, Hessisch Oldendorf, Germany |
| SafeSeal SurPhob Tips 20 µL Sterile (#VT0220) | Biozym, Hessisch Oldendorf, Germany |
| SafeSeal SurPhob Tips 100 µL Sterile (#VT0230) | Biozym, Hessisch Oldendorf, Germany |
| SafeSeal SurPhob Tips 200 µL Sterile (#VT0240) | Biozym, Hessisch Oldendorf, Germany |
| SafeSeal SurPhob Tips 1250 µL Sterile (#VT0270) | Biozym, Hessisch Oldendorf, Germany |
| Sensicare Ice Powder Free Nitrile Exam Gloves, M (#486802) | Medline, Northfield, USA |
| Serological Pipette 2 mL (#86.1252.001) | Sarstedt, Nümbrecht, Germany |
| Serological Pipette 5 mL (#86.1253.001) | Sarstedt, Nümbrecht, Germany |
| Serological Pipette 10 mL (#86.1254.001) | Sarstedt, Nümbrecht, Germany |
| Serological Pipette 25 mL (#86.1685.001) | Sarstedt, Nümbrecht, Germany |
| Supelco amber vials 12x32x4,6 mm (#27000) | Sigma-Aldrich, Saint-Louis, USA |
| Tube 5 mL (#0030119460) | Eppendorf, Hamburg, Germany |
| Culture-insert 2 well for self-insertion (#80209) | Ibidi, Gräfelfing, Germany |
| Gene Pulser/MicroPulser Electroporation Cuvettes, 0.4 cm gap (#1652081) | Bio-Rad Laboratories, Hercules, USA |

### Hardware

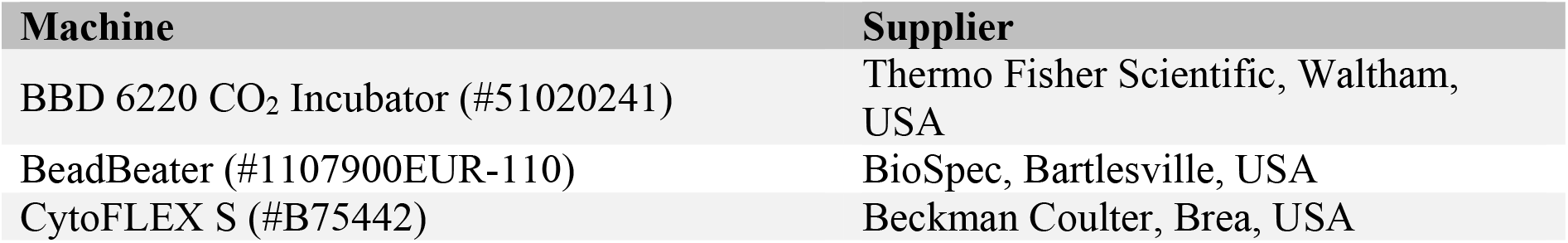
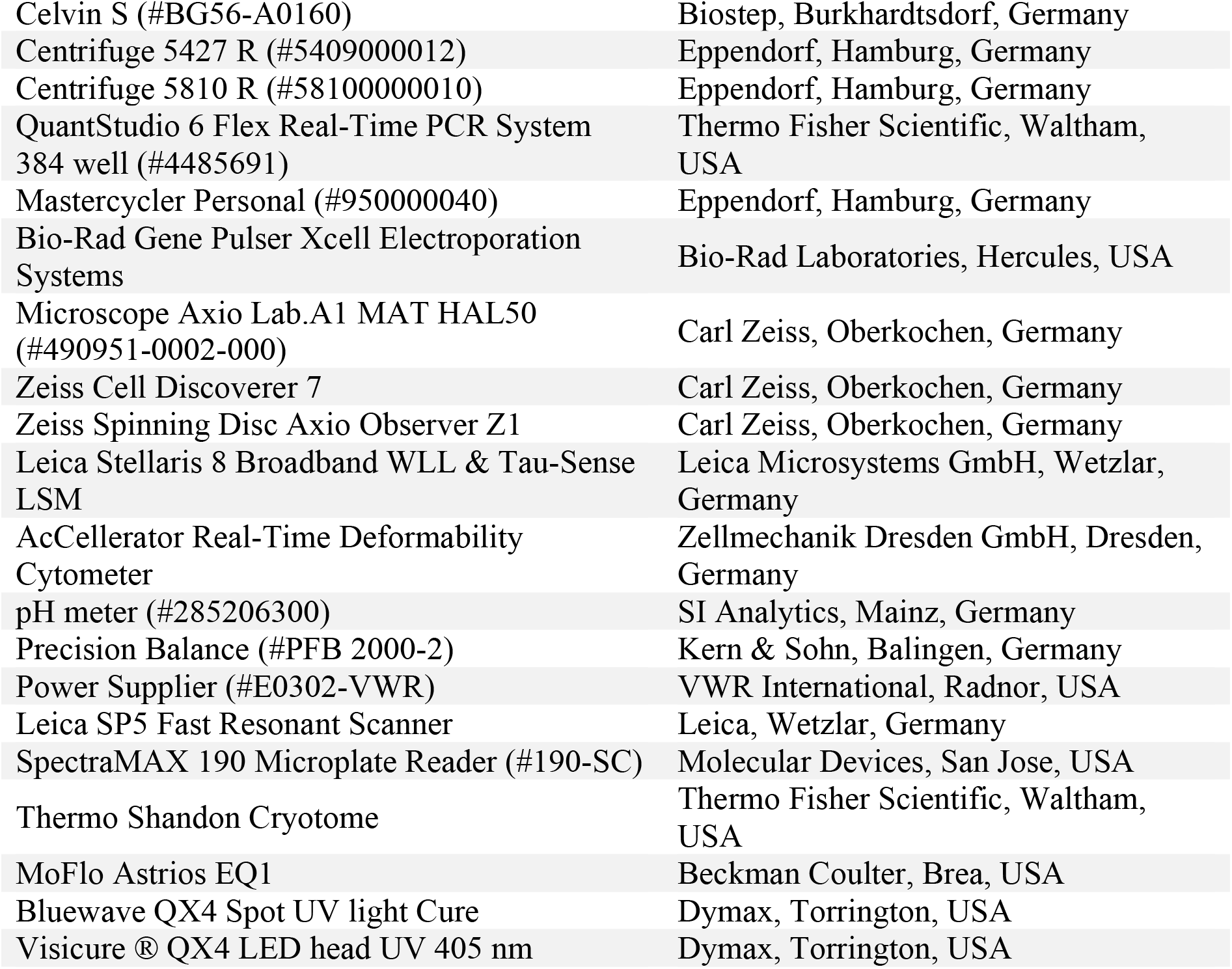

### Software

| **Software** | **Supplier** |
| --- | --- |
| Adobe Illustrator CC 2018 | Adobe, San Jose, USA |
| Adobe Photoshop CC 2018 | Adobe, San Jose, USA |
| GraphPad Prism 8 | GraphPad Software, San Diego, USA |
| Kaluza Analysis 2.1 | Beckman Coulter, Brea, USA |
| Illumina MiSeq | Illumina Inc., San Diego, USA |
| Microsoft Excel 2010 | Microsoft, Redmond, USA |
| Microsoft PowerPoint 2010 | Microsoft, Redmond, USA |
| R Software | Bell Laboratories, Murray Hill, USA |
| UPARSE | Edgar, Tiburon, USA |
| Fiji Image J v2.3.0/1.53f51 | National Institutes of Health, Bethesda, USA |
| FloJo™ software v10.8.1 | Becton Dickinson, Franklin Lakes, USA |

